# Immediate activation of chemosensory neuron gene expression by bacterial metabolites is selectively induced by distinct cyclic GMP-dependent pathways in *C. elegans*

**DOI:** 10.1101/830208

**Authors:** Jaeseok Park, Joshua D Meisel, Dennis H Kim

**Affiliations:** Division of Infectious Diseases, Boston Children’s Hospital, and Department of Pediatrics, Harvard Medical School, Boston, MA 02115, USA; Department of Biology, Massachusetts Institute of Technology, Cambridge, MA 02139, USA

**Author notes:** Department of Molecular Biology, Massachusetts General Hospital, Boston MA 02114, and Department of Genetics, Harvard Medical School, Boston, MA 02115, USA.

## Abstract

Dynamic gene expression in neurons shapes fundamental processes of the nervous systems of animals. But how different stimuli that activate the same neuron can lead to distinct transcriptional responses remains unclear. We have been studying how microbial metabolites modulate gene expression in chemosensory neurons of *Caenorhabditis elegans*. Considering the diverse environmental stimuli that can activate chemosensory neurons of *C. elegans*, we have sought to understand how specific transcriptional responses can be generated in these neurons in response to distinct cues. We have focused on the mechanism of rapid (<6 min) and selective transcriptional induction of *daf-7*, a gene encoding a TGF-β ligand that promotes bacterial lawn avoidance, in the ASJ chemosensory neurons in response to the pathogenic bacterium *Pseudomonas aeruginosa*. Here, we define the involvement of two distinct cyclic GMP (cGMP)-dependent pathways that are required for *daf-7* expression in the ASJ neuron pair in response to *P. aeruginosa*. We show that a calcium-independent pathway dependent on the cGMP-dependent protein kinase G (PKG) EGL-4, and a canonical calcium-dependent signaling pathway dependent on the activity of a cyclic nucleotide-gated channel subunit CNG-2, function in parallel to activate rapid, selective transcription of *daf-7* in response to *P. aeruginosa* metabolites. Our data suggest a requirement for PKG in promoting the fast, selective early transcription of neuronal genes in shaping responses to distinct microbial stimuli in a pair of chemosensory neurons of *C. elegans*.

**Author Summary:** The nervous systems of animals carry out the crucial roles of sensing and interpreting the external environment. When the free-living microscopic roundworm *C. elegans* is exposed to the pathogenic bacteria *Pseudomonas aeruginosa*, sensory neurons detect metabolites produced by the pathogen and induce expression of the gene for a neuroendocrine ligand called DAF-7. In turn, activity of DAF-7 is required for the full avoidance response to the *P. aeruginosa*, allowing the animals to reduce bacterial load and survive longer. Here, we systematically dissect the molecular pathway between the sensation of *P. aeruginosa* metabolites and the expression of *daf-7* in a pair of C. elegans sensory neurons. We show that the intracellular signaling molecule cyclic GMP is a key signaling intermediate. In addition, we show that there are calcium-dependent and calcium-independent pathways that are both required to engage *daf-7* expression, highlighting an organizational principle that allows neurons to distinguish between various stimuli.

## Introduction

Chemosensory systems of animals transduce external chemical stimuli into neuronal signals, with diverse roles in animal physiology (1–3). A challenge for chemosensory systems is to detect and process a wide diversity of environmental information to generate corresponding appropriate neuronal and behavioral responses. Whereas neurons utilize electrical impulses in rapid data transmission, changes in gene expression serve as a mechanism for transducing information over a longer time scale. Activity-dependent transcription of immediate-early genes has been shown to involve the activation of calcium-dependent signal transduction converging on CREB (4). Our group’s recent work has focused on understanding how changes in gene expression in chemosensory neurons of *C. elegans* can be modulated by interactions with its microbial environment (5,6).

Interactions with microbes, in a number of forms such as parasitism, symbiosis, predation, and exploitation, have shaped the evolution of animals. There has been an increasing appreciation for the role of the nervous system in recognizing and responding to microbes in the environment. Disgust, for example in response to rotting food, elicits avoidance behavior (7). At the cellular level, examples include host nociceptive neurons that have been shown to respond to microbial toxins to regulate immune responses (8), and chemosensory tuft cells, which have recently been found to sense the gut environment using canonical G-protein pathways to mediate appropriate immune responses (9–11).

The nematode *C. elegans* is usually found in rotting organic material, a complex environment in which the animal has to navigate between bacterial food, pathogens, predators, competitors, and parasites (12). With an expanded family of chemoreceptor genes in its genome and a limited set of chemosensory neurons that function to regulate diverse aspects of animal physiology, the chemosensory system of *C. elegans* enables navigation and ultimately survival in its predominantly microbial natural environment (13). We investigated how the behavior of *C. elegans* is modulated by pathogenic bacteria, specifically *Pseudomonas aeruginosa*, a devastating opportunistic pathogen of humans that is commonly found in soil and water and can also kill *C. elegans* (14).

We recently showed that the detection of virulence-associated secondary metabolites produced by *P. aeruginosa* can alter the neuronal expression pattern of *daf-7*, encoding a TGF-β ligand that regulates diverse aspects of *C. elegans* physiology (6). Specifically, whereas *daf-7* was previously known to be only expressed in the ASI head chemosensory neurons, we showed that exposure to *P. aeruginosa* causes the rapid (within six minutes) accumulation of *daf-7* mRNA in the ASJ sensory neurons (6). We also showed that exposure to the *P. aeruginosa* secondary metabolite phenazine-1-carboxamide induced an increase in calcium in the ASJ neurons. Considering that abiotic stimuli have previously been shown to increase calcium levels in the ASJ neurons (15–17), whereas induction of *daf-7* expression in the ASJ neurons is highly selective for *P. aeruginosa* metabolites, we sought to define the molecular determinants of the selective transcription of *daf-7* in the ASJ neurons in response to *P. aeruginosa*.

Here, we have taken a genetic approach to identify and characterize the signal transduction pathways in the ASJ neurons that couple the sensing of *P. aeruginosa* metabolites to the induction of *daf-7* transcription. We define distinct, parallel calcium-independent and canonical calcium-dependent pathways, each of which are dependent on cGMP signaling, which converge to selectively activate *daf-7* expression in the ASJ neurons in response to *P. aeruginosa* metabolites.

## Results

### The cyclic nucleotide-gated channel CNG-2 is required for *daf-7* expression in the ASJ neurons in response to *P. aeruginosa*

We previously described the characterization of mutants defective in the induction of *daf-7* expression in the ASJ neurons in response to *P. aeruginosa: gpa-2* and *gpa-3*, each encoding G protein alpha subunits*; tax-2* and *tax-4*, encoding components of a cyclic nucleotide-gated channel; and *daf-11*, encoding a receptor guanylate cyclase (6). These data were suggestive of a chemosensory signal transduction cascade that functions in the induction of *daf-7* expression in the ASJ neurons in response to *P. aeruginosa* metabolites.

We carried out further characterization of mutants isolated from a forward genetic screen that are defective in the induction of a *Pdaf-7::gfp* reporter transgene in the ASJ neurons in response to *P. aeruginosa*. We isolated an allele of *cng-2(qd254)* that has a splice-site mutation predicted to cause a frame-shift, rendering the protein non-functional. This mutant showed no expression of *Pdaf-7::gfp* in the ASJ neurons in the presence of *P. aeruginosa*, which we further confirmed with additional putative null alleles of *cng-2*: *tm4267, qd386* and *qd387* (Figure 1A-E, J). Unlike *tax-2* and *tax-4* mutants, *cng-2* animals did not differ from wild type in *Pdaf-7::gfp* expression in the ASI neurons (Figure 1A-E; Figure 1 ─ supplement 1A). Expression of *cng-2* cDNA specifically in the ASJ neurons using the ASJ specific promoter *trx-1* was sufficient to rescue the *Pdaf-7::gfp* expression in the ASJ neurons (Figure 1F, G, J), suggestive that CNG-2 functions in a cell-autonomous manner in the ASJ neurons to mediate *daf-7* upregulation in response to *Pseudomonas* infection.

**Figure 1.**
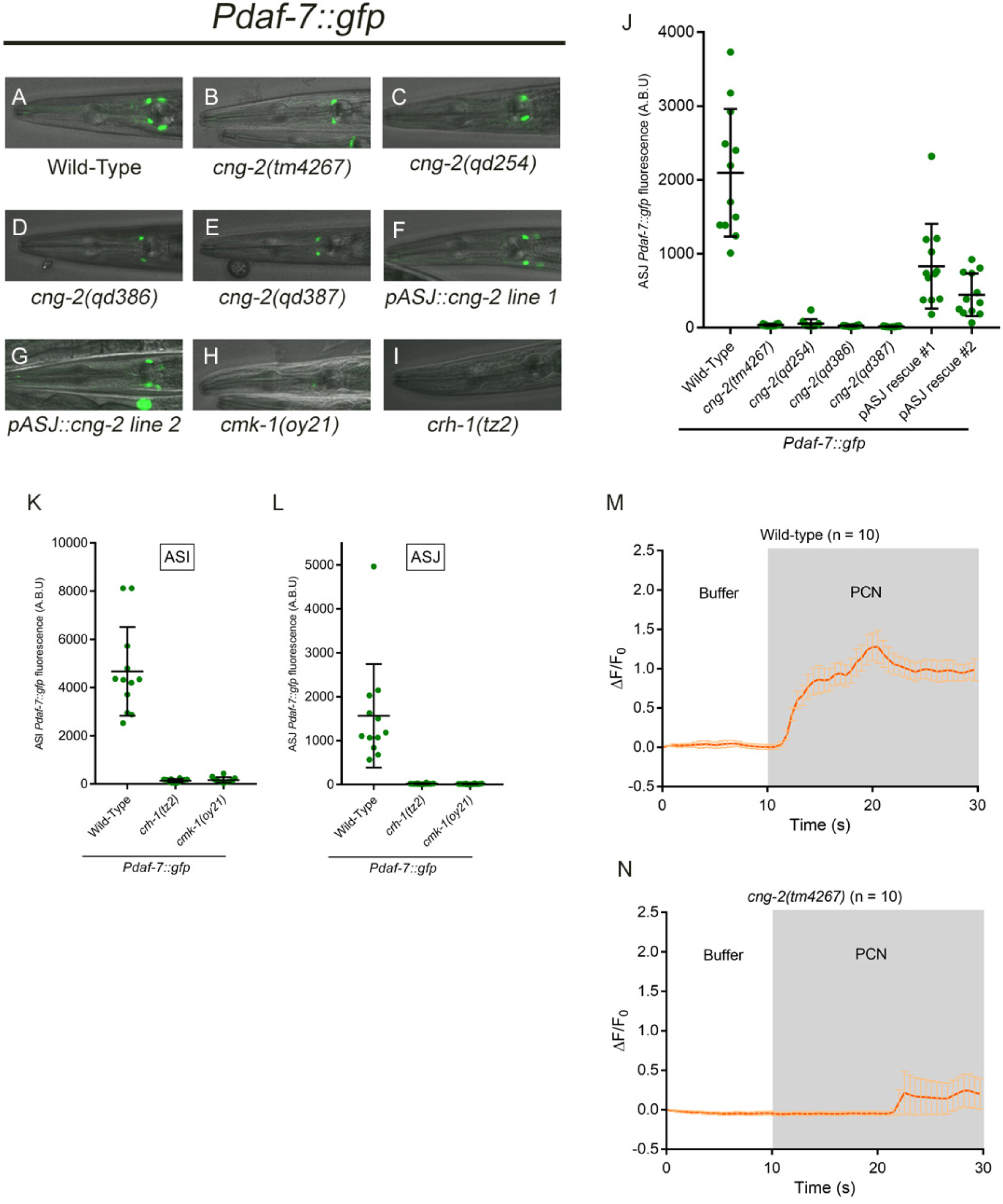
CNG-2 activates *daf-7* induction upon *P. aeruginosa* exposure in a calcium-dependent manner. All error bars indicate standard deviation. (A-I) *Pdaf-7::gfp* expression after exposure to *P. aeruginosa* for various genotypes. (J) Maximum fluorescence values of *Pdaf-7::gfp* in ASJ neurons in various *cng-2* mutant backgrounds following *P. aeruginosa* exposure. (K-L) Maximum fluorescence values of *Pdaf-7::gfp* in ASI and ASJ neurons in *crh-1* and *cmk-1* mutants following *P. aeruginosa* exposure. (M-N) GCaMP5 fluorescence change in the ASJ neurons of wild-type or *cng-2* mutant when exposed to buffer (DMSO) followed by 80 μg/ml phenazine-1-carboxamide (PCN).

We previously showed that exposure to the *P. aeruginosa* secondary metabolite, phenazine-1-carboxamide (PCN), results not only in the induction of *daf-7* expression in the ASJ neuron pair, but also in a rapid increase of calcium levels in the ASJ neurons (6). The molecular identity of CNG-2 and relevant literature on the chemosensory apparatus in the ASJ neurons (13) led us to test whether CNG-2 might be an integral part of the cation channel that is responsible for the observed calcium influx. We observed that the influx of calcium ions in the ASJ neurons that is observed upon exposure of wild-type animals to the *P. aeruginosa* metabolite phenazine-1-carboxamide was abrogated in *cng-2* animals (Figure 1M, N).

We next examined mutants carrying mutations in the genes *cmk-1*, the *C. elegans* homolog of calcium/calmodulin-dependent kinase CaMKI/IV, and *crh-1*, the *C. elegans* homolog of the transcription factor CREB, and we observed that both genes are required for *daf-7* expression in the ASJ neurons (Figure 1H, I, K, L). We note that both *cmk-1* and *crh-1* mutants also showed minimal expression of *daf-7* in the ASI neurons (Figure 1H, I, K, L), raising the possibility of additional pleiotropic effects on the development and/or physiology of the nervous systems of these mutants. Nevertheless, expressing *crh-1* cDNA in only the ASJ neurons was able to rescue *daf-7* expression in *crh-1* mutants (Figure 1-supplement 1B), indicating CRH-1 is likely to function cell-autonomously in the ASJ neurons to regulate *daf-7* expression in response to *P. aeruginosa*. These data implicate a canonical calcium-dependent signaling pathway downstream of CNG-2 that converges on CRH-1 to activate *daf-7* expression in response to phenazine-1-carboxamide.

### cGMP-dependent signal transduction activates *daf-7* expression in the ASJ neurons

The requirements for components of a cyclic nucleotide-gated channel, CNG-2/TAX-2/TAX-4, and DAF-11, a guanylate cyclase, in the induction of *Pdaf-7::gfp* expression in response to *P. aeruginosa* metabolites suggested the involvement of cGMP-dependent signaling. We demonstrated that mutations in the *gcy-12* gene, encoding another guanylate cyclase subunit, also caused markedly reduced *Pdaf-7::gfp* expression in the ASJ neurons in response to *P. aeruginosa*, but did not abolish *Pdaf-7::gfp* expression in the ASI neuron pair as was observed for the *daf-11* mutant (Figure 2A and data not shown). In addition, we observed that expression of the *gcy-12* cDNA specifically in the ASJ neurons rescued the induction of *Pdaf-7::gfp* expression in the ASJ neurons in response to *P. aeruginosa* (Figure 2A).

**Figure 2.**
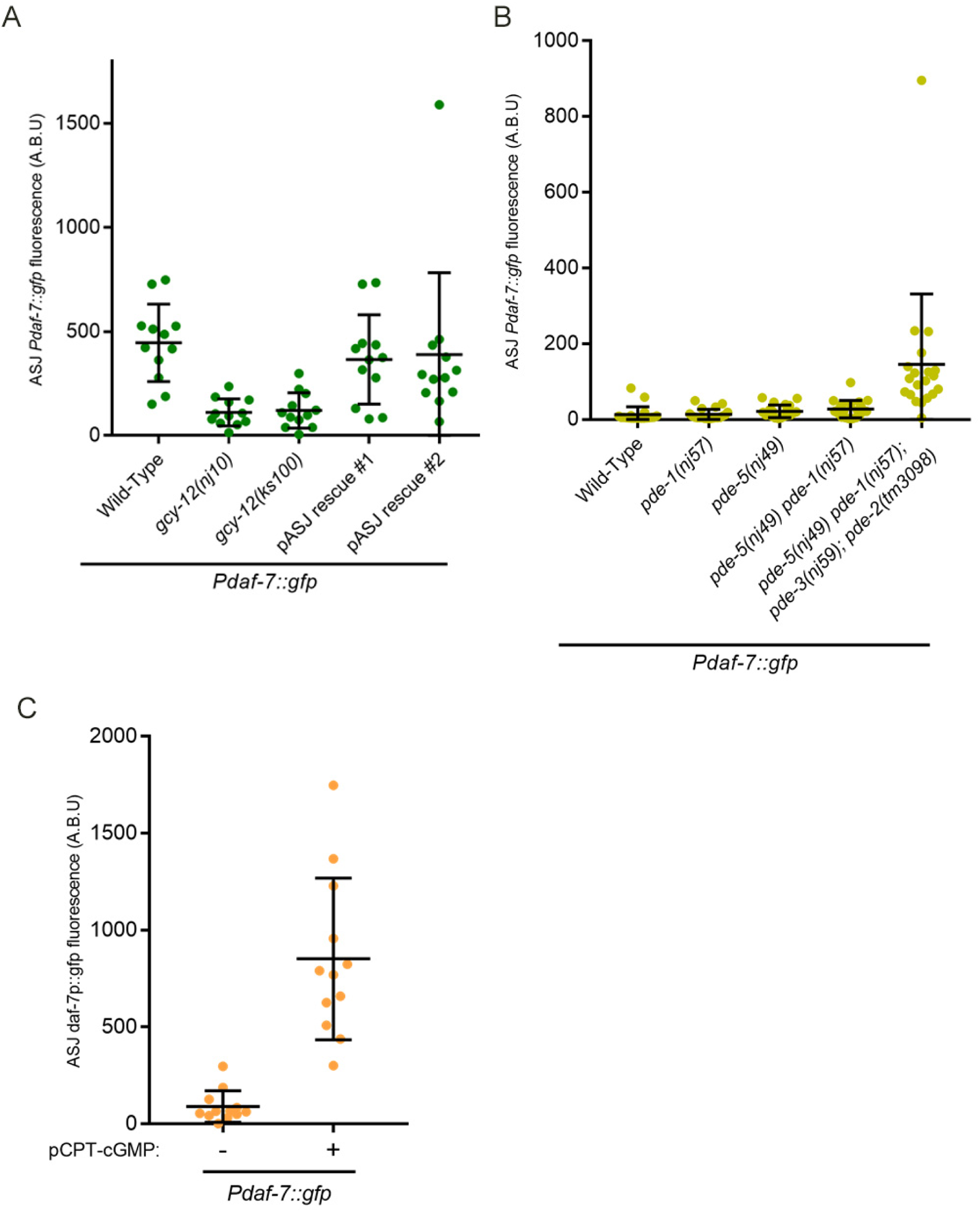
cGMP is sufficient for the induction of *daf-7* in the ASJ neurons. All error bars indicate standard deviation. (A) Maximum fluorescence values of *Pdaf-7::gfp* in ASJ neurons in various *gcy-12* mutant backgrounds following *P. aeruginosa* exposure. (B) Maximum fluorescence values of *Pdaf-7::gfp* in ASJ neurons of various phosphodiesterase (PDE) mutants. Animals were maintained on the *E. coli* strain OP50. (C) Maximum fluorescence values of *Pdaf-7::gfp* in ASJ neurons after exposure to 5 mM pCPT-cGMP. Animals were maintained on the *E. coli* strain OP50.

In order to investigate further the involvement of cGMP-dependent signaling in the induction of *daf-7* expression in the ASJ neurons, we examined the four *C. elegans* genes encoding phosphodiesterases (PDEs) that are predicted to cleave cGMP: PDE-1, PDE-2, PDE-3, and PDE-5 (18). We observed that loss-of-function of all four PDEs resulted in the induction of *Pdaf-7::gfp* expression in the ASJ neurons even in the absence of *P. aeruginosa*. Mutations in only a subset of the genes encoding PDEs conferred a considerably weaker expression of *Pdaf-7::gfp* (Figure 2B), suggestive that the PDEs function redundantly in the ASJ neurons. We also examined the effect of addition of a cell-permeable, non-hydrolysable analog of cGMP, pCPT-cGMP, to wild-type animals in the absence of *P. aeruginosa*, and we observed the marked induction of expression of *daf-7* in the ASJ neurons (Figure 2C).

### The cGMP-dependent protein kinase G EGL-4 upregulates *daf-7* expression in ASJ neurons in response to *P. aeruginosa*

The involvement of cGMP- and calcium-dependent signaling support a role for canonical activity-dependent signaling pathways in the induction of *daf-7* expression in response to *P. aeruginosa* metabolites. However, prior studies have shown that multiple stimuli including low pH, *E. coli* supernatant, sodium chloride, temperature changes, and even water can cause calcium influx in the ASJ neurons without the robust upregulation of *daf-7* observed in the presence of *P. aeruginosa* (15–17). We sought to define additional, calcium-independent mechanisms that might be involved in the selective transcriptional response to *P. aeruginosa* metabolites.

The dependence of *daf-7* expression in the ASJ neurons on cGMP led us to consider the involvement of the cGMP-dependent protein kinase G (PKG), EGL-4. EGL-4 has been implicated in various phenotypes including egg-laying behavior, chemosensory behavior, sleep-like state, satiety signaling, and aversive learning behaviors (19–29). We observed that presumptive loss-of-function *egl-4(n478)* and *egl-4(n479)* mutants exhibited a lack of *Pdaf-7::gfp* expression in the ASJ neurons in response to *P. aeruginosa* (Figure 3A-C, E). Expression of *egl-4* cDNA in the ASJ neurons was sufficient to rescue *daf-7* expression on *P. aeruginosa* (Figure 3D, E). A gain-of-function allele, *egl-4(ad450)*, exhibited detectable expression of *daf-7* expression in the ASJ neurons even in the absence of *P. aeruginosa* (Figure 4A). In order to examine whether EGL-4 functioned in a calcium-dependent or calcium-independent manner, we examined how *egl-4* loss-of-function affected the influx of calcium into the ASJ neurons observed upon exposure to phenazine-1-carboxamide. We found that unlike *cng-2(tm4267)* mutants, *egl-4(n479)* mutants showed a wild-type calcium level increase in ASJ neurons upon exposure to phenazine-1-carboxamide (Figure 3F, G).

**Figure 3.**
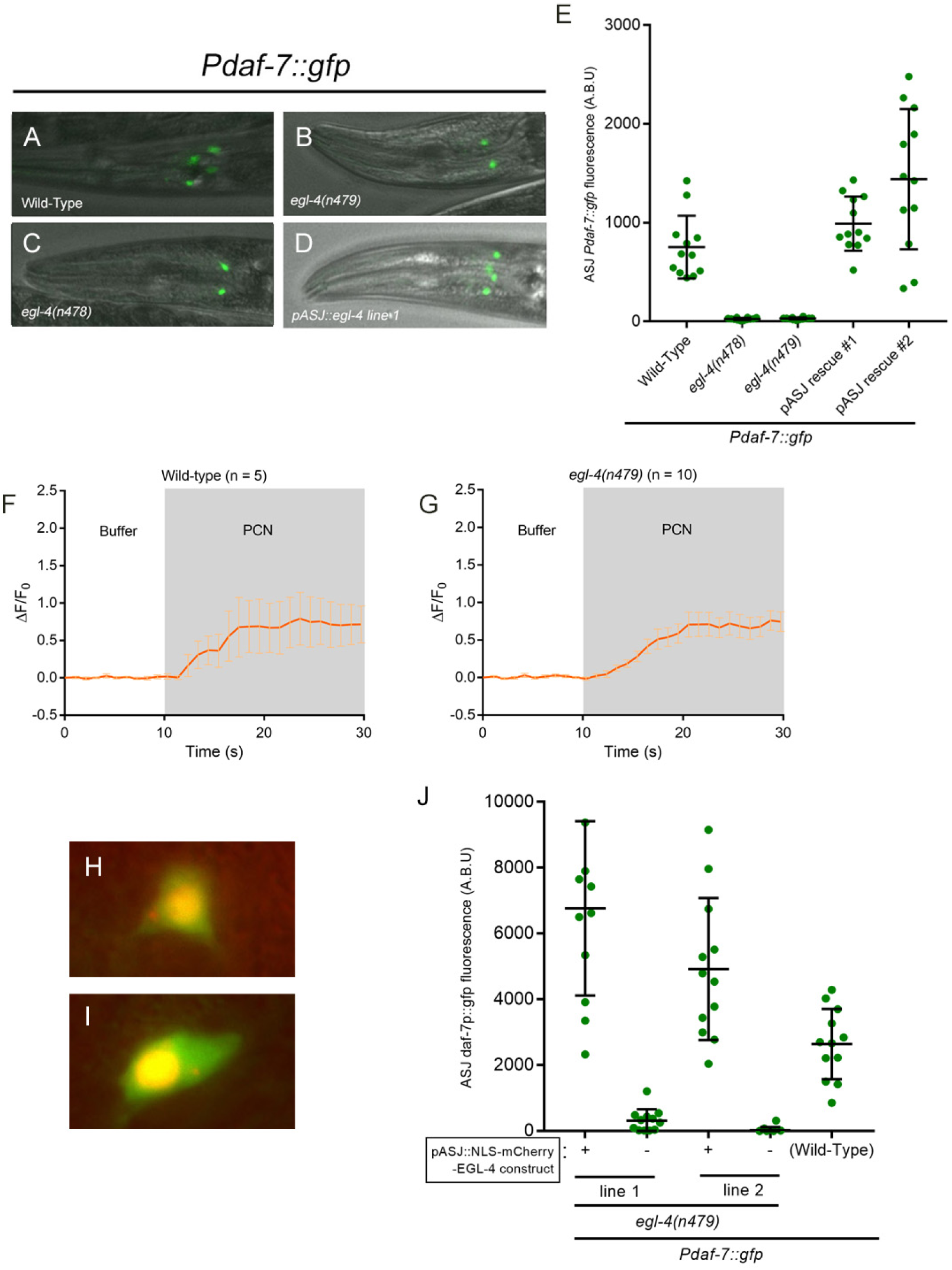
*egl-4* mutants show defects in *daf-7* induction on *P. aeruginosa*. All error bars indicate standard deviation. (A-D) *Pdaf-7::gfp* expression after exposure to *P. aeruginosa* for various *egl-4* backgrounds. (E) Maximum fluorescence values of *Pdaf-7::gfp* in ASJ neurons in various *egl-4* mutant backgrounds following *P. aeruginosa* exposure. (F-G) GCaMP5 fluorescence change in the ASJ neurons when exposed to buffer (DMSO) followed by 66 μg/ml phenazine-1-carboxamide (PCN) in wild-type or *egl-4* mutants. (H-I) NLS-mCherry-EGL-4 proteins are localized to the nucleus. GFP is observed throughout the ASJ neurons, outlining the cells. (J) Maximum fluorescence values of *Pdaf-7::gfp* in ASJ neurons in *egl-4(n479)* mutants containing the NLS-mCherry-EGL-4 constructs. Imaging followed exposure to *P. aeruginosa.* Sibling populations with or without the construct were compared in each line.

**Figure 4.**
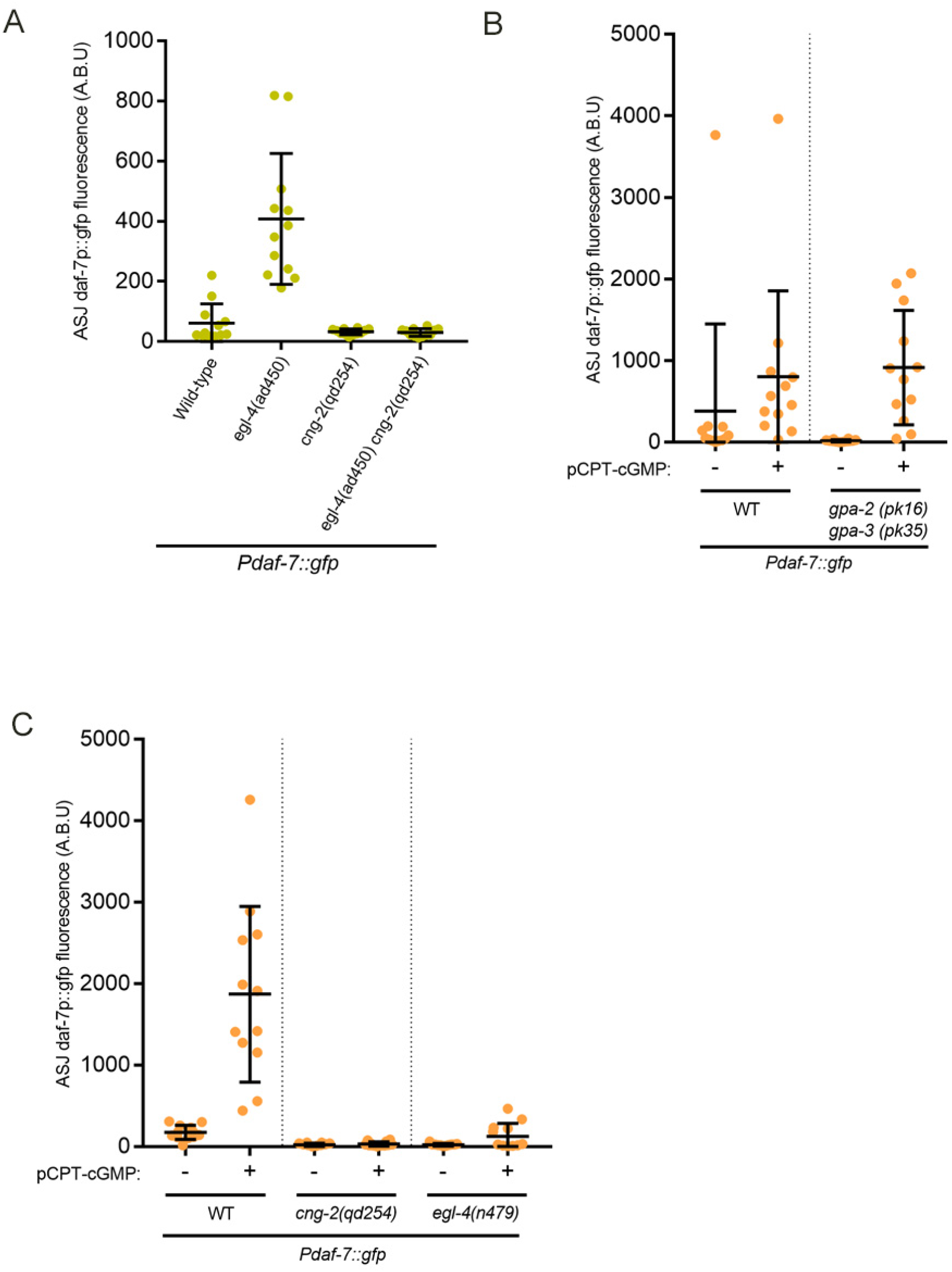
EGL-4 and CNG-2 work concurrently in parallel pathways downstream of cGMP to induce *daf-7*. All error bars indicate standard deviation. (A) Epistasis analysis using *cng-2*(loss-of-function) and *egl-4*(gain-of-function) alleles. Animals were maintained on the *E. coli* strain OP50. (B-C) Maximum fluorescence values of *Pdaf-7::gfp* in ASJ neurons of various mutants following exposure to 5mM pCPT-cGMP. Animals were maintained on the *E. coli* strain OP50.

PKGs have been reported to function in the nucleus to mediate gene expression in some cases (20,30). To investigate the possibility that EGL-4 functions in the nucleus to mediate its effects in ASJ neurons to regulate *daf-7* expression, we expressed mCherry::EGL-4 with an additional nuclear localization sequence (NLS) at the N-terminus, resulting in an EGL-4-derived transgene that was localized to the nucleus (Figure 3H, I). We observed that expression of this nuclear EGL-4 construct rescued the *daf-7* expression defect in the ASJ neurons (Figure 3J). These data are consistent with a role for EGL-4 in the nucleus to regulate *daf-7* transcription in the ASJ neurons in response to *P. aeruginosa*.

To better define the respective roles and interaction between *cng-2* and *egl-4*, we again utilized the gain-of-function allele of *egl-4*, *ad450*. In the *egl-4(ad450) cng-2(qd254)* double mutant, the expression we observed in the *egl-4(ad450)* mutant was abolished (Figure 4A), which suggested that basal CNG-2-dependent calcium-dependent signaling in the absence of *P. aeruginosa* is required for the observed *daf-7* expression. We further sought to gain clarity regarding the pathway involving *cng-2* and *egl-4* by seeing how each of the mutants might change their *daf-7* expression in response to the addition of pCPT-cGMP. We first tested mutants of the heterotrimeric G-protein *gpa-2(pk16)* and *gpa-3(pk35)*. These proteins are thought to act in the initial steps of the chemosensory cascade by associating with the presumptive receptor for *P. aeruginosa* metabolites, and we have previously shown that the *gpa-2 gpa-3* double mutant is defective in *daf-7* transcription in response *to P. aeruginosa* (6). Consistent with this prediction, adding pCPT-cGMP to *gpa-2 gpa-3* double mutants elicited the same induction of *Pdaf-7::gfp* expression in the ASJ neurons as observed in wild-type animals (Figure 4B). However, when pCPT-cGMP was added to *cng-2* and *egl-4* mutants, the response was markedly attenuated or absent (Figure 4C), consistent with roles for CNG-2 and EGL-4 functioning downstream of and dependent on a cGMP signal in the induction of *daf-*7 expression in the ASJ neuron pair in response to *P. aeruginosa* metabolites.

## Discussion

The neuroendocrine TGF-beta ligand, DAF-7, is rapidly transcribed in the ASJ neurons upon exposure to *P. aeruginosa* metabolites (6). In this study, we have identified and characterized several mutants that define the signaling mechanisms coupling the sensing of *P. aeruginosa* metabolites to the induction of *daf-7* expression in the ASJ neuron pair. The identification of cGMP-dependent signaling proteins, CNG-2 and EGL-4, pointed to a pivotal role for cGMP. This was further corroborated by the involvement of guanylate cyclases, GCY-12 and DAF-11, the effect of inactivating multiple redundant phosphodiesterases acting on cGMP, and chemical induction of *Pdaf-7::gfp* expression in the ASJ neurons using pCPT-cGMP. Our data suggest a model for the sequence of cellular signaling events that are initiated by the detection of *P. aeruginosa* metabolites and result in the rapid induction of *daf-7* expression (Figure 5). In particular, cGMP-dependent signaling through CNG-2 activates a canonical calcium-dependent signaling pathway that likely activates CREB. In parallel, cGMP-dependent signaling activates EGL-4, which functions in the nucleus in concert with calcium-dependent signaling converging on CREB. Both pathways are required for the full activation of *daf-7*, as inactivation of either pathways alone resulted in the inability to robustly upregulate *daf-7*.

**Figure 5.**
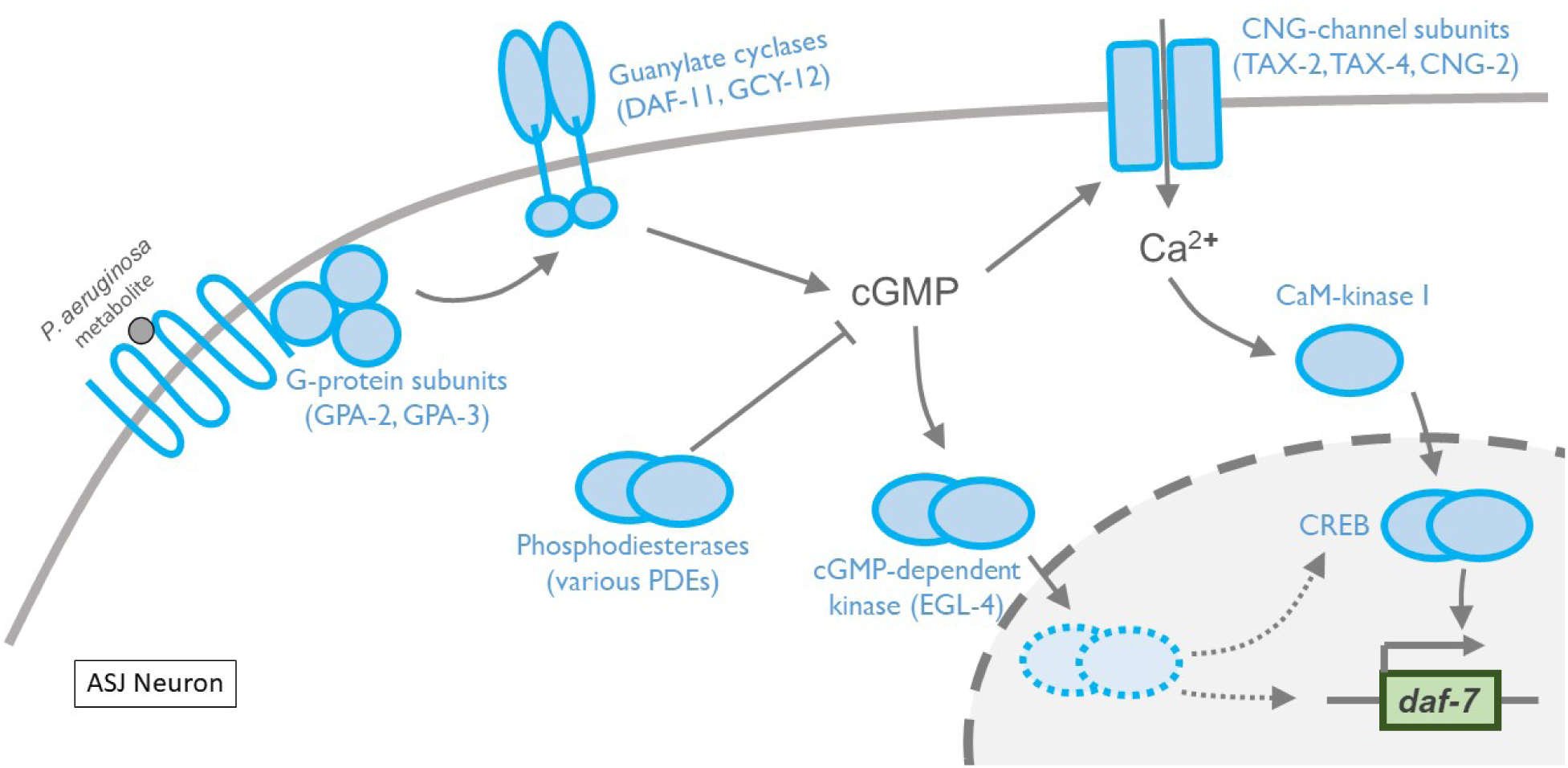
Fast transcription of *daf-7* is selectively induced by activation of calcium-dependent and calcium-independent pathways in ASJ neurons. A schematic describing the current model for the sensory transduction pathway in the ASJ neurons resulting in fast neuronal gene transcription in response to *P. aeruginosa* metabolite phenazine-1-carboxamide (PCN). The model highlights the role of canonical signal transduction pathway molecules as well the added role of the cGMP-dependent kinase EGL-4 as one of the two parallel pathways required for the induction of *daf-7*. Note that the activation of both pathways are required for the full induction of *daf-7*.

TAX-2, TAX-4, and CNG-2 are subunits for the hetrotetrameric cyclic nucleotide-gated channel, and DAF-11 and GCY-12 are subunits for the dimeric guanylyl cyclase. In *tax-2, tax-4*, and *daf-11* mutants, *daf-7* expression is lost in both the ASI and ASJ neurons. In contrast, *daf-7* expression in ASI neurons of *cng-2* and *gcy-12* mutants is relatively intact, even as *daf-7* expression in ASJ neurons is lost, suggestive of more ASJ-specific roles for CNG-2 and GCY-12. Neuron-specific activities of CNG-2 and GCY-12 may confer distinct biochemical properties to the CNG channels and guanylyl cyclases in different neurons. Such organization would be consistent with what is seen in other organisms: for example, the CNG channels in the rods and cones of the mammalian retina have different subunit compositions and thus have different biochemical properties differentially optimized for the functions of each cell type (31).

Our data point to a key role for calcium-independent signaling through EGL-4 in the selective transcriptional activation of *daf-7* in ASJ neurons. Various external stimuli have been shown to activate ASJ neurons as measured by calcium level changes, such as low pH, salt, and *E. coli* supernatant (17), whereas the robust expression of *daf-7* in the ASJ neurons is activated selectively by *P. aeruginosa*. Loss-of-function *egl-4* mutants are unable to induce *daf-7* in ASJ neurons on *P. aeruginosa* (Figure 3E), and the *daf-*7 transcriptional response to the cGMP analog pCPT-cGMP is severely compromised (Figure 4C), underlining the requirement of EGL-4 in *daf-7* expression. Moreover, *egl-4* mutants have wild-type calcium influx in the ASJ neurons when exposed to phenazine-1-carboxamide, implying the necessity, but not sufficiency of calcium influx in the induction of *daf-7* expression in the ASJ neurons in response to *P. aeruginosa*. Thus, EGL-4 and activation and calcium influx seem to work together to regulate *daf-7* expression. Calcium and cGMP have been known to work collaboratively to regulate immediate early gene expression in various neuron types (32–35). However, our data uniquely demonstrates a key role for cGMP-dependent signaling functioning in concert with canonical calcium-dependent signaling pathways in a pair of primary sensory neurons, activated by physiological environmental ligands.

*C. elegans* is anatomically restricted in its neuronal system, with only 302 neurons to carry out sensation, data processing, and motor output all at once. Such constraints dictate that unlike mammalian olfactory neurons, *C. elegans* chemosensory neurons may have to process multiple types of stimuli in a single neuron, while retaining the ability to distinguish between them. The ASJ neurons routinely use calcium-dependent signaling to mediate signal transduction to a wide variety of stimuli, but our data suggest that select stimuli such as secondary pathogen metabolites can be distinguished and linked to gene transcription by engaging calcium-independent PKGs in addition to calcium-dependent signals (Figure 5). Whereas how PKGs can be activated selectively in response to different stimuli remains to be explored further, our data provide an indication of how transcriptional responses in sensory neurons of *C. elegans* may be gated through distinct signal transduction pathways to result in selective changes in gene expression to promote adaptive behaviors.

## Material and Methods

### *C. elegans* Strains

All animals were maintained and fed as previously described (36). The animals were incubated at 20°C unless any of the strains were considered temperature-sensitive, in which case they were grown at 16°C. Please see Supplementary Methods for a complete list of strains used in this study.

### *Pdaf-7::gfp* induction assays and quantification

For experiments quantifying the level of *Pdaf-7::gfp* on the *Pseudomonas aeruginosa* strain PA14, bacteria was cultured overnight in 3 mL LB broth at 37°C, and the following day 7 μl was seeded onto 3.5cm slow-killing assay (SKA) plates as described previously (14). The seeded plates were maintained at 37°C overnight and then transferred to room temperature, where they were kept additional two days before experiments. To preemptively rid animals of bacterial contamination, gravids were bleached to get a large amount of eggs. Animals were loaded onto PA14 at stage L4 and then were kept at 25°C for 14-16 hours before quantification. For assays using pCPT-cGMP, pCPT-cGMP was added to SKA plates in mixed DMSO and water, with the resulting concentration in plates to be 5 mM. Plates were left overnight for the chemical to diffuse. The next day, 5 μl inoculate of *E. coli* strain OP50 was seeded to the middle, and plates were kept in room temperature overnight before experiments commenced. Animals were similarly egg-prepped for this condition as noted above. L4s were loaded onto the center of the SKA plates and kept at 20°C for 17-20 hrs before quantification.

### Quantification of *Pdaf-7p::gfp* levels

Animals were mounted on glass slides with agarose pads and anesthetized with 50 mM sodium azide. Animals were imaged using a Zeiss Axioimager Z1. Quantification of GFP brightness was conducted with FIJI by obtaining maximum fluorescence values within the ASJ, or ASI neurons. Y-axes are denoted by arbitrary brightness units (A.B.U.).

### Generation of transgenic animals

The *trx-1* promoter (1.1 kb) was amplified by PCR from genomic DNA (37), and *unc-54* 3’ UTR was amplified from Fire Vector pPD95.75. *cng-2* cDNA generously provided by P Sengupta, *gcy-12* cDNA by M. Fujiwara, *egl-4* cDNA by N. D. L’Etoile, and *crh-1* cDNA by C. T. Murphy, were all respectively amplified by PCR. Finally, the *trx-1* promoter, respective cDNAs, and the *unc-54* 3’ UTR were cloned into plasmids using NEBuilder® HiFi DNA Assembly (New England Biolabs, Ipswich, MA). The plasmids were microinjected at 40 - 50 ng/μl concentration, along with *ofm-1p::gfp* as a co-injection market at 30 - 40 ng/μl for ASJ specific expression. For generation of strains with calcium indicators, amplified *trx-1* promoter was fused with GCaMP5G to express the indicator in ASJ neurons only.

### Calcium imaging

The animals were immobilized and exposed to soluble compound in a controlled manner using a microfluidics chip as previously described (38). Imaging was carried out at 40x with a Zeiss Axiovert S100 inverted microscope equipped with an Andor iXon EMCCD camera. Stimulus was given at noted concentrations. Phenazine-1-carboxamide was obtained from Princeton BioMolecular Research (Princeton, NJ). Data analysis was done with a custom MATLAB script written by Nikhil Bhatla.

## Acknowledgements

We thank members of the Kim lab, members of the Horvitz lab, and Yun Zhang for advice and discussions. We thank Na An and H. Robert Horvitz, Manabi Fujiwara, Noelle D. L’Etoile, Coleen T. Murphy, and Piali Sengupta for strains and reagents. We thank the *Caenorhabditis* Genetics Center (supported by National Institutes of Health - Office of Research Infrastructure Programs, P40 OD010440) and the National BioResource Project for providing strains. We thank Kurt Broderick at MIT Microsystems Technology Laboratories for assistance with microfluidics chip fabrication. This work was supported by National Institutes of Health grant GM084477 (to D.H.K).

---

**Figure 1 – figure supplement 1.**
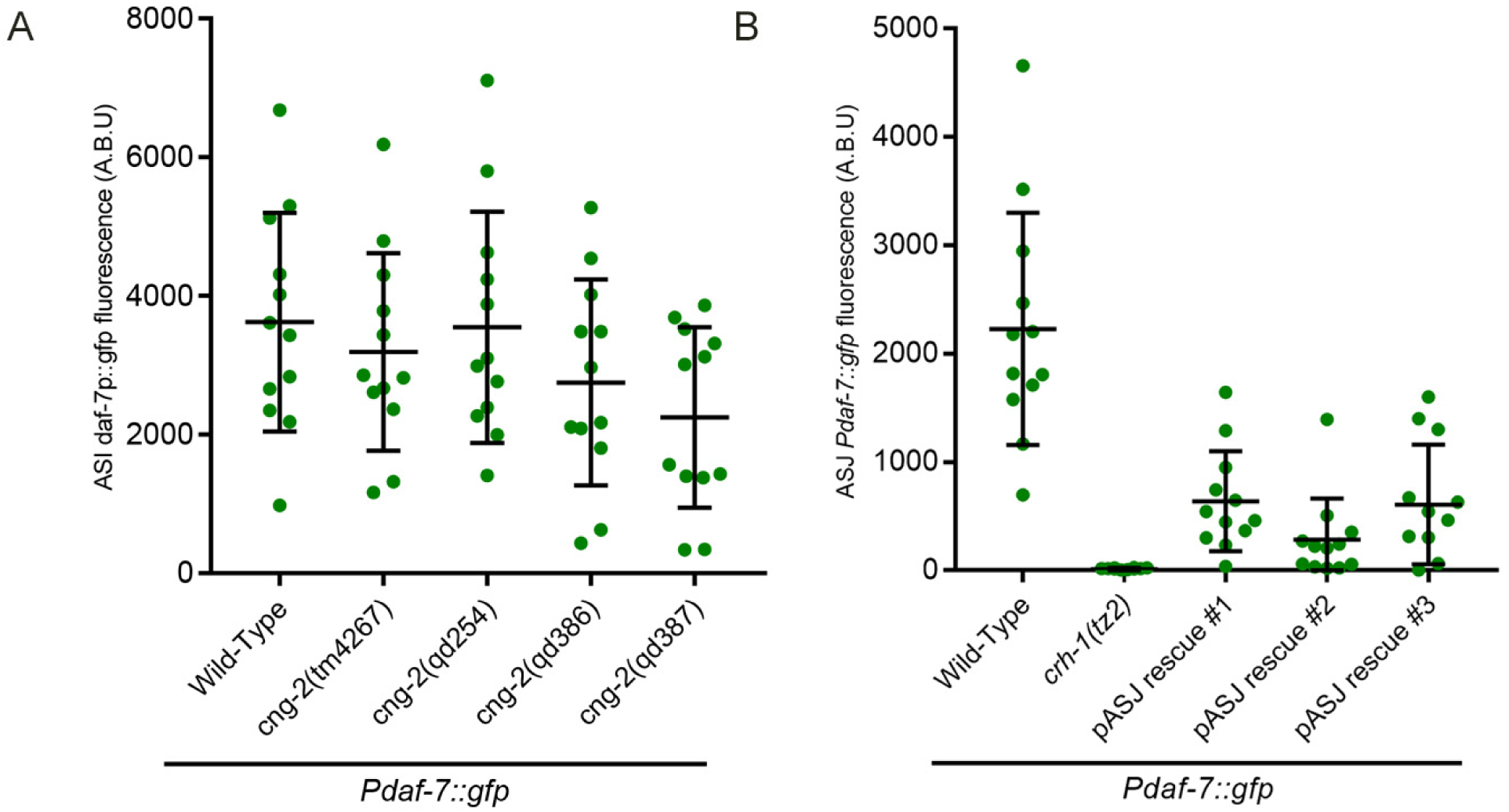
(A) *Pdaf-7::gfp* levels of ASI neurons in *cng-2* mutants after exposure to *P. aeruginosa*. (B) Expressing *crh-1* cDNA specifically in the ASJ neurons restores *daf-7* expression on *P. aeruginosa*.

## Supplemental Methods

**Table S1.** All strains used in this study

| STRAIN | GENOTYPE |
| --- | --- |
| N2 | Wild-Type |
| FK181 | *ksIs2[Pdaf-7::GFP + rol-6(su1006)]* |
| ZD727 | *ksIs2; crh-1(tz2)* |
| ZD899* | *ksIs2; cng-2(tm4267)* |
| ZD882 | *ksIs2; cng-2(qd254)* |
| ZD887 | *ksIs2; gpa-2(pk16) gpa-3(pk35)* |
| ZD1048 | *ksIs2; cmk-1(oy21)* |
| ZD1401 | *ksIs2; gcy-12(nj10)* |
| ZD1922 | *ksIs2; egl-4(n478); him-5(e1490)* |
| ZD1737 | *qdEx136[trx-1p::GCaMP5; ofm-1::gfp]* |
| ZD1949 | *ksIs2; egl-4(ad450); him-5(e1490)* |
| ZD2020 | *cng-2(tm4267); qdEx136[trx-1p::GCaMP5; ofm-1p::gfp]* |
| ZD2090 | *ksIs2; gcy-12(ks100)* |
| ZD2091 | *pde-5(nj49) ksIs2* |
| ZD2105 | *pde-5(nj49) pde-1(nj57) ksIs2* |
| ZD2150 | *pde-1(nj57) ksIs2* |
| ZD2151 | *pde-5(nj49) pde-1(nj57) ksIs2; pde-3(nj59); pde-2(tm3098)* |
| ZD2170 | *egl-4(n479); qdEx136[trx-1p::GCaMP5::unc-54 5'UTR; ofm-1p::gfp]* |
| ZD2534 | *ksIs2; cng-2(qd386)* |
| ZD2535 | *ksIs2; cng-2(qd387)* |
| ZD2250 | *ksIs2; cng-2(qd254); qdEx[pJP3(trx-1p::cng-2cDNA::unc-54UTR) + ofm-1p::gfp] Line #1* |
| ZD2251 | *ksIs2; cng-2(qd254); qdEx[pJP3(trx-1p::cng-2cDNA::unc-54UTR) + ofm-1p::gfp] Line #2* |
| ZD2275 | *ksIs2; egl-4(n479ts)* |
| ZD2283 | *ksIs2; egl-4(ad450) cng-2(qd254)* |
| ZD2307 | *ksIs2; egl-4(n479); qdEx[trx-1p::mCherry::egl-4cDNA + ofm-1p::gfp] Line #1* |
| ZD2308 | *ksIs2; egl-4(n479); qdEx[trx-1p::mCherry::egl-4cDNA + ofm-1p::gfp] Line #2* |
| ZD2526 | *ksIs2; gcy-12(ks100); qdEx[trx-1p::gcy-12cDNA::unc-54UTR + ofm-1p::gfp] Line #1* |
| ZD2527 | *ksIs2; gcy-12(ks100); qdEx[trx-1p::gcy-12cDNA::unc-54UTR + ofm-1p::gfp] Line #2* |
| ZD2536 | *ksIs2; egl-4(n479); qdEx[pJP17(trx-1p::SV40-nls::mCherry::egl-4cDNA::unc54UTR) + ofm-1p::gfp] Line #1* |
| ZD2537 | *ksIs2; egl-4(n479); qdEx[pJP17(trx-1p::SV40-nls::mCherry::egl-4cDNA::unc54UTR) + ofm-1p::gfp] Line #2* |
| ZD2538 | *ksIs2; egl-4(n479); qdEx[pJP17(trx-1p::SV40-nls::mCherry::egl-4cDNA::unc54UTR) + ofm-1p::gfp] Line #3* |
| ZD2545 | *ksIs2; crh-1(tz2); qdEx[pJP13(trx-1p::crh-1cDNA::unc-54UTR) + ofm-1p::gfp]*<br>Line #1 |
| ZD2546 | *ksIs2; crh-1(tz2); qdEx[pJP13(trx-1p::crh-1cDNA::unc-54UTR) + ofm-1p::gfp]*<br>Line #2 |
| ZD2547 | *ksIs2; crh-1(tz2); qdEx[pJP13(trx-1p::crh-1cDNA::unc-54UTR) + ofm-1p::gfp]*<br>Line #3 |

\**tm4267* is currently classified as lethal or sterile by the National BioResource Project of Japan and was originally maintained on a balancer. However, *cng-2(tm4267)* is viable, as we were able to backcross away the linked lethal mutation.

## References

1. Hildebrand JG, Shepherd GM. Mechanisms of olfactory discrimination: converging evidence for common principles across phyla. Annu Rev Neurosci. 1997;20:595–631.

2. Yarmolinsky DA, Zuker CS, Ryba NJP. Common sense about taste: from mammals to insects. Cell. 2009 Oct 16;139(2):234–44.

3. Yohe LR, Brand P. Evolutionary ecology of chemosensation and its role in sensory drive. Curr Zool. 2018 Aug;64(4):525–33.

4. Flavell SW, Greenberg ME. Signaling mechanisms linking neuronal activity to gene expression and plasticity of the nervous system. Annu Rev Neurosci. 2008;31:563–90.

5. Hilbert ZA, Kim DH. Sexually dimorphic control of gene expression in sensory neurons regulates decision-making behavior in C. elegans. Elife. 2017 24;6.

6. Meisel JD, Panda O, Mahanti P, Schroeder FC, Kim DH. Chemosensation of bacterial secondary metabolites modulates neuroendocrine signaling and behavior of C. elegans. Cell. 2014;159(2):267–80.

7. Curtis V, Biran A. Dirt, disgust, and disease. Is hygiene in our genes? Perspect Biol Med. 2001;44(1):17–31.

8. Chiu IM, Heesters BA, Ghasemlou N, Von Hehn CA, Zhao F, Tran J, et al. Bacteria activate sensory neurons that modulate pain and inflammation. Nature. 2013 Sep 5;501(7465):52–7.

9. Gerbe F, Sidot E, Smyth DJ, Ohmoto M, Matsumoto I, Dardalhon V, et al. Intestinal epithelial tuft cells initiate type 2 mucosal immunity to helminth parasites. Nature. 2016 Jan 14;529(7585):226–30.

10. Howitt MR, Lavoie S, Michaud M, Blum AM, Tran SV, Weinstock JV, et al. Tuft cells, taste-chemosensory cells, orchestrate parasite type 2 immunity in the gut. Science. 2016 Mar 18;351(6279):1329–33.

11. von Moltke J, Ji M, Liang H-E, Locksley RM. Tuft-cell-derived IL-25 regulates an intestinal ILC2-epithelial response circuit. Nature. 2016 Jan 14;529(7585):221–5.

12. Schulenburg H, Félix M-A. The Natural Biotic Environment of Caenorhabditis elegans. Genetics. 2017;206(1):55–86.

13. Bargmann C. Chemosensation in C. elegans. Wormbook. 2006;1–29.

14. Tan MW, Mahajan-Miklos S, Ausubel FM. Killing of Caenorhabditis elegans by Pseudomonas aeruginosa used to model mammalian bacterial pathogenesis. Proc Natl Acad Sci USA. 1999 Jan 19;96(2):715–20.

15. Ohta A, Ujisawa T, Sonoda S, Kuhara A. Light and pheromone-sensing neurons regulates cold habituation through insulin signalling in *Caenorhabditis elegans*. Nature Communications. 2014 Jul 22;5:4412.

16. Wang W, Qin L-W, Wu T-H, Ge C-L, Wu Y-Q, Zhang Q, et al. cGMP Signalling Mediates Water Sensation (Hydrosensation) and Hydrotaxis in *Caenorhabditis elegans*. Scientific Reports. 2016 Feb 19;6:19779.

17. Zaslaver A, Liani I, Shtangel O, Ginzburg S, Yee L, Sternberg PW. Hierarchical sparse coding in the sensory system of Caenorhabditis elegans. Proc Natl Acad Sci. 2015;112(4):1185–9.

18. Liu J, Ward A, Gao J, Dong Y, Nishio N, Inada H, et al. C. elegans phototransduction requires a G protein-dependent cGMP pathway and a taste receptor homolog. Nat Neurosci. 2010 Jun;13(6):715–22.

19. Cho CE, Brueggemann C, L’Etoile ND, Bargmann CI. Parallel encoding of sensory history and behavioral preference during Caenorhabditis elegans olfactory learning. Elife. 2016 06;5.

20. Juang B-T, Gu C, Starnes L, Palladino F, Goga A, Kennedy S, et al. Endogenous nuclear RNAi mediates behavioral adaptation to odor. Cell. 2013 Aug 29;154(5):1010–22.

21. Krzyzanowski MC, Brueggemann C, Ezak MJ, Wood JF, Michaels KL, Jackson CA, et al. The C. elegans cGMP-dependent protein kinase EGL-4 regulates nociceptive behavioral sensitivity. PLoS Genet. 2013;9(7):e1003619.

22. Lee JI, M O Damien, Jeffery E-A, Juang B-T, Kaye JA, Hamilton SO, et al. Nuclear entry of a cGMP-dependent kinase converts transient into long-lasting olfactory adaptation. Proc National Acad Sci. 2010;107(13):6016–21.

23. L’Etoile ND, Coburn CM, Eastham J, Kistler A, Gallegos G, Bargmann CI. The Cyclic GMP-Dependent Protein Kinase EGL-4 Regulates Olfactory Adaptation in C. elegans. Neuron. 2002;36(6):1079–89.

24. Halloran DM, Hamilton OS, Lee JI, Gallegos M, L’Etoile ND. Changes in cGMP levels affect the localization of EGL-4 in AWC in Caenorhabditis elegans. PLoS ONE. 2012;7(2):e31614.

25. O’Halloran DM, Altshuler-Keylin S, Lee JI, L’Etoile ND. Regulators of AWC-mediated olfactory plasticity in Caenorhabditis elegans. PLoS Genet. 2009 Dec;5(12):e1000761.

26. Raizen DM, Zimmerman JE, Maycock MH, Ta UD, You Y, Sundaram MV, et al. Lethargus is a Caenorhabditis elegans sleep-like state. Nature. 2008;451(7178):nature06535.

27. Trent C, Tsuing N, Horvitz HR. Egg-laying defective mutants of the nematode Caenorhabditis elegans. Genetics. 1983 Aug;104(4):619–47.

28. van der Linden AM, Wiener S, You YJ, Kim K, Avery L, Sengupta P. The EGL-4 PKG acts with KIN-29 salt-inducible kinase and protein kinase A to regulate chemoreceptor gene expression and sensory behaviors in Caenorhabditis elegans. Genetics. 2008;180(3):1475–91.

29. You Y, Kim J, Raizen DM, Avery L. Insulin, cGMP, and TGF-beta signals regulate food intake and quiescence in C. elegans: a model for satiety. Cell Metab. 2008 Mar;7(3):249–57.

30. Gudi T, Lohmann S, Pilz R. Regulation of gene expression by cyclic GMP-dependent protein kinase requires nuclear translocation of the kinase: identification of a nuclear localization signal. Mol Cell Biol. 1997;17(9):5244–54.

31. Podda MV, Grassi C. New perspectives in cyclic nucleotide-mediated functions in the CNS: the emerging role of cyclic nucleotide-gated (CNG) channels. Pflugers Arch. 2014 Jul;466(7):1241–57.

32. Belsham DD, Wetsel WC, Mellon PL. NMDA and nitric oxide act through the cGMP signal transduction pathway to repress hypothalamic gonadotropin-releasing hormone gene expression. EMBO J. 1996 Feb 1;15(3):538–47.

33. Chen Y, Zhuang S, Cassenaer S, Casteel DE, Gudi T, Boss GR, et al. Synergism between calcium and cyclic GMP in cyclic AMP response element-dependent transcriptional regulation requires cooperation between CREB and C/EBP-beta. Mol Cell Biol. 2003 Jun;23(12):4066–82.

34. Lee SA, Park JK, Kang EK, Bae HR, Bae KW, Park HT. Calmodulin-dependent activation of p38 and p42/44 mitogen-activated protein kinases contributes to c-fos expression by calcium in PC12 cells: modulation by nitric oxide. Brain Res Mol Brain Res. 2000 Jan 10;75(1):16–24.

35. Peunova N, Enikolopov G. Amplification of calcium-induced gene transcription by nitric oxide in neuronal cells. Nature. 1993 Jul 29;364(6436):450–3.

36. Brenner S. The genetics of Caenorhabditis elegans. Genetics. 1974;77(1):71–94.

37. Fierro-González JC, Cornils A, Alcedo J, Miranda-Vizuete A, Swoboda P. The thioredoxin TRX-1 modulates the function of the insulin-like neuropeptide DAF-28 during dauer formation in Caenorhabditis elegans. PLoS ONE. 2011 Jan 27;6(1):e16561.

38. Chronis N, Zimmer M, Bargmann CI. Microfluidics for in vivo imaging of neuronal and behavioral activity in Caenorhabditis elegans. Nat Methods. 2007;4(9):727–31.

